# Evaluation of Host Defense Peptide (CaD23)-Antibiotic Interaction and Mechanism of Action: Insights from Experimental and Molecular Dynamics Simulations Studies

**DOI:** 10.1101/2021.06.26.450050

**Authors:** Darren Shu Jeng Ting, Jianguo Li, Chandra S. Verma, Eunice T. L. Goh, Mario Nubile, Leonardo Mastropasqua, Dalia G. Said, Roger W. Beuerman, Rajamani Lakshminarayanan, Imran Mohammed, Harminder S. Dua

**Author notes:** Equally contributing authors (Imran Mohammed, Harminder S. Dua). **Corresponding author:** Professor Harminder S. Dua, **Address:** Academic Ophthalmology, Division of Clinical Neuroscience, School of Medicine, University of Nottingham, Nottingham, UK. and 62025.

## Abstract

**Background/aim:** Host defense peptides (HDPs) have the potential to provide a novel solution to antimicrobial resistance (AMR) in view of their unique and broad-spectrum antimicrobial activities. We had recently developed a novel hybrid HDP based on LL-37 and human beta-defensin-2, named CaD23, which was shown to exhibit good *in vivo* antimicrobial efficacy against *Staphylococcus aureus* in a bacterial keratitis murine model. This study aimed to examine the potential CaD23-antibiotic synergism and to evaluate the underlying mechanism of action of CaD23.

**Methods:** Antimicrobial efficacy was determined using minimum inhibitory concentration (MIC) assay with broth microdilution method. Peptide-antibiotic interaction was evaluated against *S. aureus*, methicillin-resistant *S. aureus* (MRSA), and *Pseudomonas aeruginosa* using established checkerboard assay and time-kill kinetics assay. Fractional inhibitory concentration index (FICI) was calculated and interpreted as synergistic (FICI<0.5), additive (FICI between 0.5-1.0), indifferent (FICI between >1.0 and ≤4), or antagonistic (FICI>4). SYTOX green uptake assay was performed to determine the membrane-permeabilising action of CaD23. Molecular dynamics (MD) simulations were performed to evaluate the interaction of CaD23 with bacterial and mammalian mimetic membranes.

**Results:** CaD23-amikacin and CaD23-levofloxacin combination treatment exhibited a strong additive effect against *S. aureus* SH1000 (FICI=0.56) and MRSA43300 (FICI=0.56) but a borderline additive-to-indifferent effect against *P. aeruginosa* (FIC=1.0-2.0). CaD23 (at 25 μg/ml; 2x MIC) was able to achieve complete killing of *S. aureus* within 30 mins. When used at sub-MIC concentration (3.1 μg/ml; 0.25x MIC), it was able to expedite the antimicrobial action of amikacin against *S. aureus* by 50%. The rapid antimicrobial action of CaD23 was attributed to the underlying membrane-permeabilising mechanism of action, evidenced by the SYTOX green uptake assay and MD simulations studies. MD simulations revealed that cationicity, alpha-helicity, amphiphilicity and hydrophobicity (related to the Trp residue at C-terminal) play important roles in the antimicrobial action of CaD23.

**Conclusions:** CaD23 is a novel membrane-active synthetic HDP that can enhance and expedite the antimicrobial action of antibiotics against Gram-positive bacteria when used in combination. MD simulation serves as a useful tool in dissecting the mechanism of action and guiding the design and optimisation of HDPs.

## INTRODUCTION

Antimicrobial resistance (AMR) is currently one of the major global health threats.^1,2^ By 2050, it is estimated to cause 10 million deaths and cost the global economy up to 100 trillion USD if the issue remains untackled.^3^ In addition, non-systemic infections, including ocular and skin infections, are being increasingly affected by drug-resistant pathogens, which usually result in poor prognosis.^2,4,5^ In view of the colossal impact on global health and economy, various initiatives and strategies have been proposed and implemented to tackle AMR. These include establishment of antimicrobial stewardship to monitor the use of antimicrobial agents and the rise of AMR, development of new drugs and vaccines, drug repurposing, and incentivising pharmaceutical companies for investing in antimicrobial drug development.^2^

Infectious keratitis (IK) represents the 5^th^ leading cause of blindness globally.^4^ It can be caused by a wide range of organisms, including bacteria, fungi, viruses, and parasites, particularly Acanthamoeba.^6–13^ Broad-spectrum topical antibiotic treatment is the current mainstay of treatment for IK, but the management is being challenged by the low culture yield,^4,6^ polymicrobial infection,^8,14,15^ and emerging AMR.^16–19^ In addition, adjuvant procedures / surgeries such as therapeutic cross-linking,^20^ amniotic membrane transplantation,^21^ and therapeutic / tectonic keratoplasty^22^ are often required to manage uncontrolled infection and its complications, including corneal melting and perforation. All these issues highlight the need for new treatment for IK.

Host defense peptides (HDPs) have shown promise as a novel solution to AMR in view of their unique and broad-spectrum antimicrobial activities.^23^ These HDPs are usually highly cationic and amphiphilic, with ~30-50% hydrophobicity.^24–26^ The cationic amino acid residues facilitate the binding of HDPs onto the anionic bacterial membrane (via electrostatic interactions), while the hydrophobic residues interact with the lipid tail region of the membrane, culminating in membrane disruption, leakage of cytoplasmic contents and subsequent cell death.^27^ In addition to the direct antimicrobial activity, HDPs exhibit anti-biofilm, anti-tumour, immunomodulatory, chemotactic and wound-healing properties, offering a wide range of potential therapeutic applications.^23,28^

However, several barriers, including cytotoxicity to host cells and stability in the host / infective environment, have so far hindered the clinical translation of HDP-based antimicrobial therapy.^29^ To overcome these barriers, some research groups have explored the use of peptide-antibiotic combination therapy as a means to exploit the peptide-antibiotic synergistic effect for treating various types of infections.^30–34^ This attractive antimicrobial strategy not only helps extend the lifespan and broaden the antibacterial spectrum of conventional antibiotics, but also reduces the dose-dependent toxicity associated with HDPs and antibiotics.^35^

Recently, our group had demonstrated that CaD23, a hybrid derivative of human cathelicidin (LL-37) and human beta-defensin (HBD)-2, exhibited a more rapid *in vitro* antimicrobial action than conventional antibiotics such as amikacin.^36^ However, the mechanism of action has not been fully elucidated. In addition, while CaD23 exhibited reasonable *in vivo* efficacy at a concentration of 0.05% (500 μg/ml), the use of a higher concentration of CaD23 to achieve stronger antimicrobial effect was prohibited by the toxicity, as observed in the wound healing study. Therefore, to overcome this limitation, we aimed to examine the potential synergism / interaction between CaD23 and commonly used antibiotics for IK, including levofloxacin and amikacin.^37^ In addition, we aimed to determine the mechanism of action of CaD23 using a combination of experimental and molecular dynamics (MD) simulations studies.

## MATERIALS AND METHODS

### Chemicals and antibiotics

All the peptides were commercially produced by Mimotopes (Mimotopes Pty. Ltd., Mulgrave Victoria, Australia) *via* traditional solid phase Fmoc synthesis method. All the synthetic peptides were purified by reverse-phase high performance liquid chromatography (RP-HPLC) to >95% purity and characterised by mass spectrometry. In view of the hydrophobicity, CaD23 (sequence: KRIVQRIKDWLRKLCKKW) was first fully dissolved in 50 μl of dimethyl sulfoxide (DMSO) followed by dilution in sterile, de-ionised water to achieve a final concentration of 1 mg/ml peptide in 0.5% v/v DMSO. Further dilution was performed for specific assays as required. All the assays described in this study were conducted in biological duplicate and in at least two independent experiments, with appropriate positive controls (PCs) and negative controls (NCs). Antibiotics, including levofloxacin and amikacin, were purchased from Sigma-Aldrich (Merck Life Science UK Ltd., Dorset, UK).

### Types of microorganisms used

A range of Gram-positive and Gram-negative bacteria were used in this study. These included laboratory-strain methicillin-sensitive *Staphylococcus aureus* (SH1000 and ATCC SA29213), methicillin-resistant *S. aureus* (ATCC MRSA43300), *Pseudomonas aeruginosa* ATCC PA19660 (cytotoxic strain), and *P. aeruginosa* ATCC PA27853 (invasive strain). Both cytotoxic and invasive *P. aeruginosa* strains were used in the experiments as previous studies had demonstrated the difference in virulence.^38,39^

### Determination of antimicrobial efficacy

*In vitro* antimicrobial efficacy of CaD23 and the antibiotics was determined using the established minimum inhibitory concentration (MIC) assay with broth microdilution method approved by the Clinical and Laboratory Standards Institute (CLSI).^40^ Briefly, the microorganisms were cultured on Tryptone Soya Agar (TSA) and incubated overnight for 18-21 hours at 37°C. Bacterial inoculums were subsequently prepared using the direct colony suspension method.^40^ Three to five bacterial colonies were obtained from the agar plate and inoculated into an Eppendorf tube containing 1 ml of cation-adjusted Muller-Hinton broth (caMHB, Merck), consisting of 20-25 mg/L calcium ions (Ca^2+^) and 10-12.5 mg/L magnesium ions (Mg^2+^). The bacterial suspension was adjusted to achieve a turbidity equivalent to 0.1 OD_600_ or 0.5 MacFarland, containing ~1.5 × 10^8^ colony-forming unit (CFU)/ml, which was then further diluted in 1:150 in caMHB to reach a final bacterial concentration of ~1×10^6^ colony forming units (CFU)/ml. Subsequently, 50 μl of 1×10^6^ CFU/ml bacteria and 50 μl of treatment / controls were added into each well for the MIC assay. The MIC values, defined as the lowest concentration of the antimicrobial agent that prevented any visible growth of bacteria, were determined after 18-21 hours of incubation at 37°C.

### Determination of the peptide-antibiotic interaction

The peptide-antibiotic interaction was determined using two methods, namely the checkerboard assay and the time-kill kinetics assay.

#### Checkerboard assay

The peptide-antibiotic synergism was examined using the established checkerboard assay described in the previous study.^32^ A 96-well polypropylene plate (Plate A) was used to prepare 8 replicate horizontal rows of CaD23 in twofold serial dilutions [from 400 μg/ml (1^st^ column) to 6.25 μg/ml (7^th^ column), and caMHB in the last (8^th^) column; final volume of 25 μl per well]. Another 96-well polystyrene plate (Plate B) was used to prepare 8 replicate vertical columns of an antibiotic, either amikacin (an aminoglycoside) or levofloxacin (a fluoroquinolone), in twofold serial dilutions [from 20 μg/ml (1^st^ row) to 0.313 μg/ml (7^th^ row), and 0 μg/ml in the last (8^th^) row; final volume of 30 μl per well]. Subsequently, 25 μl of antibiotic from each well of Plate B was transferred to the corresponding wells of Plate A (1:1 ratio of peptide and antibiotic). The bacterial suspension was prepared as above and 50 μl of 1×10^6^ CFU/ml bacteria was added into each well (1:1 ratio of treatment and bacteria; final concentration of 5 × 10^5^ CFU/ml bacteria per well). The final concentration of CaD23 in each row was 100 μg/ml (1^st^ column) to 1.56 μg/ml (7^th^ column) and the final concentration of antibiotic in each column was 5 μg/ml (1^st^ row) to 0.078 μg/ml (7^th^ row). Growth control and sterility control were included in each experiment. The MIC was calculated as above after 18-21 hours of incubation with treatment at 37°C.

The fractional inhibitory concentration index (FICI) is calculated using the formula: (MIC_CaD23(combined)_ / MIC_CaD23(alone)_) + (MIC_antibiotic(combined)_ / MIC_antibiotic(alone)_) and was interpreted as synergistic (FICI <0.5), additive (FICI between 0.5-1.0), indifferent (FICI between >1.0 and ≤4), or antagonistic (FICI >4).

#### Time-kill kinetics assay

Time-kill kinetics assay was performed to determine the time and concentration-dependent antimicrobial activity of CaD23 and amikacin against SH1000. The bacterial suspension (with a concentration of 1 × 10^6^ CFU/ml) was prepared using the similar method as described in the MIC assay. 50 μl of bacteria was then incubated with 50 μl of respective treatment, consisting of either CaD23 alone, amikacin alone, or combined CaD23-amikacin. Bacterial suspension incubated with sterile de-ionised water (dH_2_O) in 1:1 ratio was used as the growth control. At 0 min, 15 min, 30 min, 1 hour, 2 hour, 4 hours, and 24 hours, 10 μl of the treatment / bacteria mixture was removed from each well and was serially diluted (1:10 dilution) in sterile phosphate buffer solution (PBS). The diluted suspension (20 μl) was subsequently removed and plated on Muller-Hinton agar (MHA) in duplicate for bacterial counting after incubation for 18-21 hours at 37°C.

### Evaluation of the mechanism of action

#### SYTOX green uptake assay

SYTOX green is a membrane-impermeable dye that activates and fluoresces upon binding to the DNA. The assay was performed using a previously established method, with a slight modification.^41^ Briefly, the bacteria were cultured overnight in MHB (20 μl) for 16-18 hours. Subsequently, the bacterial suspension was vortexed, washed twice and suspended in sterile HEPES buffer solution (5 mM HEPES, 5 mM glucose, 7.4 pH) to obtain an OD_600_ of 0.3. An aliquot of 5 mM SYTOX green stock solution in DMSO was added to the bacterial suspension to obtain a final dye concentration of 2 μM. The mixture was incubated for 15 minutes at room temperature while being protected from light. The dye-loaded cell suspension (600 μl) was then added into a stirring quartz cuvette and inserted into a QuantaMaster spectrofluorometer for fluorescence time-based scan at 504 nm excitation and 523 nm emission. Once a constant fluorescence level was achieved, a concentrated peptide solution in water (1 μl) was added into the cuvette in order to obtain a desired final concentration of CaD23 (2x MIC) in the cell suspension. The change in fluorescence intensity was monitored until a stable range was observed. Maximum fluorescence was documented via the addition of Triton-X (final concentration of Triton-X 0.1% (v/v) in 600 μl cell suspension) into the cuvette. The fluorescence intensity (I) of the peptide-treated suspension was calculated and plotted as: (I_peptide_ / I_Triton-X(max)_) × 100%

#### Molecular dynamics simulations

Established molecular dynamics (MD) simulations-based models were used to examine the interactions between the synthetic peptides and models of the bacterial and mammalian membranes, using the GROMACS 5.1 package.^16^ The ability of peptide to permeate or interact with the bacterial membrane and mammalian membrane served as a proxy for its antimicrobial efficacy and toxicity, respectively. The bacterial membrane was modelled using a mixture of phosphoethanolamine and phosphatidylglycerol lipids (3:1 ratio) whilst the mammalian membrane was modelled using phosphotidylcholine. Each membrane patch consists of 128 lipid molecules. The peptide was modelled using the AMBER14sb force field, and the lipid molecules were modelled using the AMBER lipid17 force field. Initially, the peptide, modelled in a helical conformation, was placed 4 nm above the membrane center, followed by solvation with water molecules using the TIP3 model of each system.^42^ Counter ions were added to neutralise each system. Each system was first subjected to 500 steps of energy minimisation, followed by 20 ps of MD simulation in the canonical NVT ensemble (N = constant number; V = volume; T = temperature). Each system was first simulated for 400 ns to allow the peptide to adsorb on the membrane surface. Due to the complex free energy landscape of the peptide-membrane system, the time scale required to reach the equilibrium state was considerably lengthy. To overcome this difficulty, 400 ns simulations of simulated annealing, as outlined by Farrotti et al.,^43^ were performed. In each simulated annealing cycle, the temperature of the system was increased from 300 K to 375 K in 50 ps steps, followed by a 1 ns simulation at 300 K. This was followed by 400 ns of normal MD simulation at 300 K. The LINCS algorithm^44^ was applied to restrain the bond between hydrogen atoms and heavy atoms, enabling a time step of 2 fs. Both Lennard-Jones and short-range electrostatic interactions were set to extend to 0.9 nm, while the long range electrostatic interactions were calculated using particle mesh Ewald method.^45^ The temperature and pressure were controlled by Nose–Hoover^46^ and semi-isotropic Parrinello–Rahman algorithms,^47^ respectively.

## RESULTS

### Peptide-antibiotic interaction

#### Checkerboard assay

The MICs of CaD23, amikacin and levofloxacin against Gram-positive and Gram-negative bacteria are presented in **Table 1**. A number of peptide-antibiotic combinations were examined for their interactive antimicrobial effect against both Gram-positive and Gram-negative bacteria (**Table 2**). It was found that both CaD23-amikacin and CaD23-levofloxacin combinations achieved strong additive effects against SH1000 (FICI = 0.563) and MRSA43300 (FICI = 0.563). On the other hand, the effect of CaD23-amikacin was indifferent against PA19660 (FICI = 1.08) and borderline additive against PA27853 (FICI = 1.0), whereas the effect of CaD23-levofloxacin against PA19660 and PA27853 was indifferent (both FICI = 2.0).

#### Time-kill kinetics assay

Based on the results of the checkerboard assay, the concentration- and time-dependent antimicrobial effect of combined CaD23-amikacin against SH1000 was further explored. The MIC of CaD23 and amikacin against SH1000 was 12.5 μg/ml and 1.25 μg/ml, respectively. When CaD23 was used alone at the concentration of 25 μg/ml (2x MIC), it was able to achieve 99.9% and 100% killing of SH1000 by 15 mins and 30 mins post-treatment, respectively. This was significantly faster than amikacin at 10 μg/ml (4x MIC) or 25 μg/ml (10x MIC), which both achieved 99.9% and 100% killing of SH1000 by 2 hours and 4 hours post-treatment (i.e. 8 times slower). The addition of CaD23 (3.1 μg/ml; 0.25x MIC) expedited the antimicrobial action of amikacin (10 μg/ml) against SH1000 by 4 times (for 99.9% killing) and 2 times (for 100% killing). In addition, combined CaD23 (3.1 μg/ml; 0.25x MIC) and amikacin (2.5 μg/ml; 2x MIC) was able to achieve 99.9% and 100% killing by 1 hour and 2 hours, respectively. This was 2 times faster than amikacin when used alone at 10 μg/ml or 25 μg/ml, suggesting that combination treatment enables a more effective killing and lowers the treatment concentration required for effective killing.

### Mechanism of action of CaD23

#### SYTOX green uptake assay

SYTOX green uptake assay was performed to study the underlying mechanism of action of CaD23 against *S. aureus* ATCC SA29213 (MIC = 25 μg/ml). It was shown that CaD23 at 50 μg/ml (2x MIC) exhibited rapid membrane permeabilisation of SA29213, with 60% SYTOX green uptake observed within seconds of treatment and reaching 80% membrane permeabilisation at around 8 mins post-treatment (**Figure 2**).

**Figure 1.**
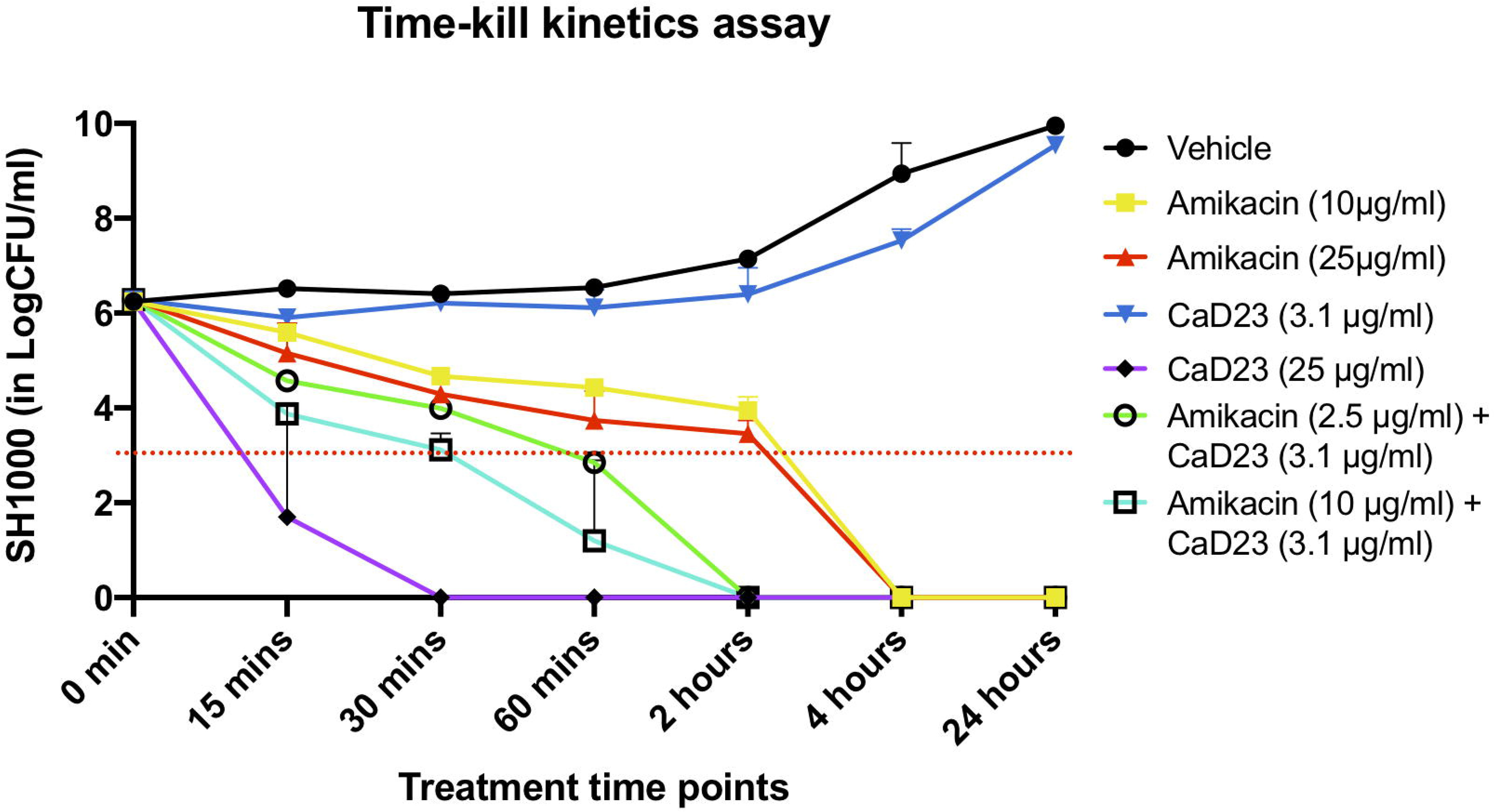
Time-kill kinetics assay examining the time- and concentration-dependent anti-bacterial effect of CaD23 (0.25x MIC and 2x MIC), amikacin (8x and 20x MIC) and combined CaD23-amikacin against *S. aureus* (SH1000) over 24 hours. The MIC of CaD23 and amikacin against SH1000 was 12.5 μg/ml and 1.25 μg/ml. SH1000 incubated with phosphate buffer solution (PBS) serves as the untreated control. “0 min” represents the starting inoculum, which is around 6 log_10_ CFU/ml. The red dotted horizontal line at 3 log_10_ CFU/ml signifies the threshold of significant bacterial killing (defined as 99.9% or 3 log_10_ CFU/ml reduction of the bacterial viability compared to the starting inoculum). Data is presented as mean ± standard deviation (depicted in error bars) of two independent experiments performed in biological duplicate. MIC = Minimum inhibitory concentration; CFU = Colony-forming unit

**Figure 2.**
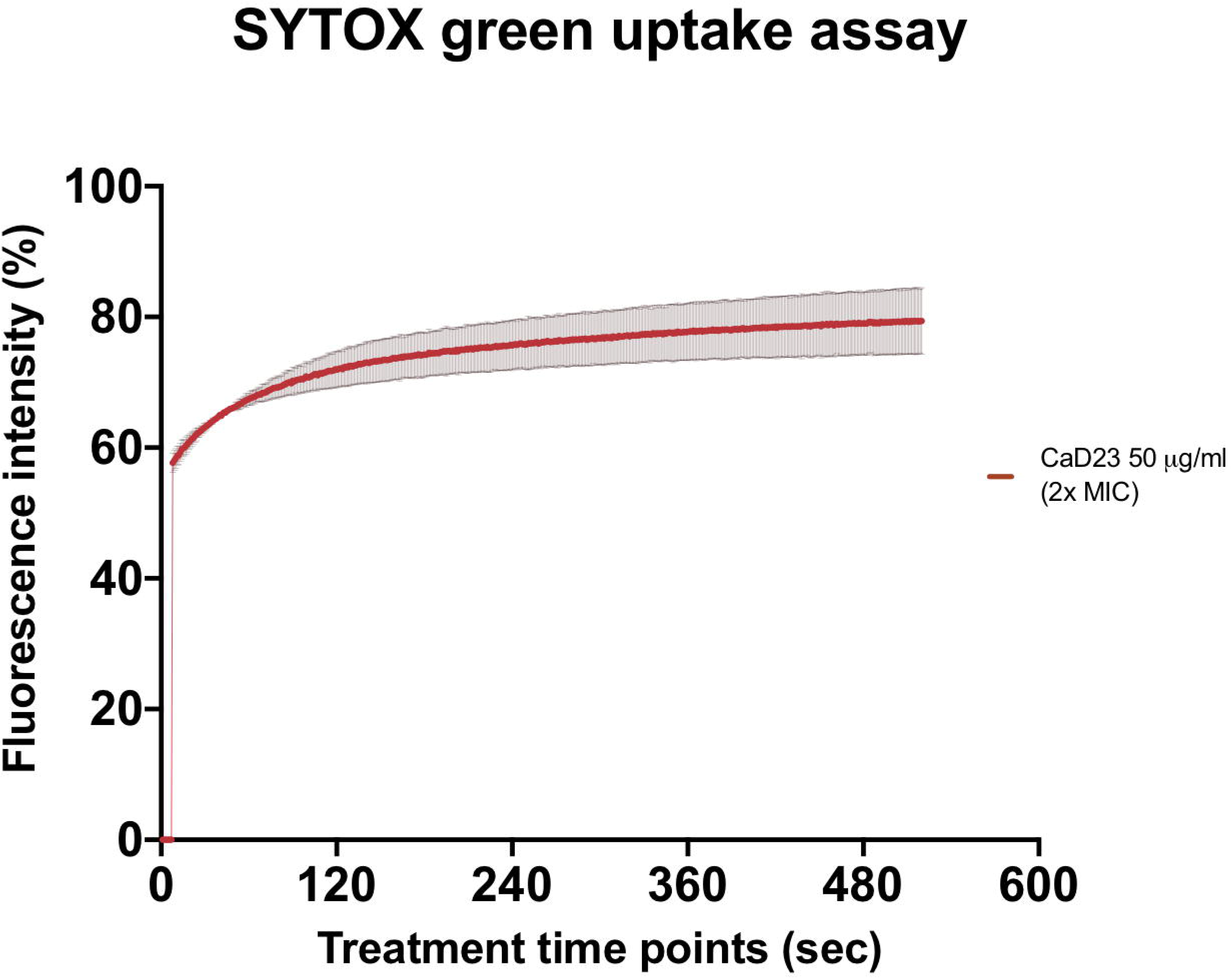
Membrane permeabilising action of CaD23 against *S. aureus* ATCC SA29213 determined by SYTOX green uptake assay. The graph demonstrating rapid membrane permeabilising action of CaD23 (50 μg/ml; 2x MIC) against SA29123, with a 60% increase in fluorescence intensity (due to SYTOX green uptake) within seconds of treatment and plateaued at ~80% fluorescence intensity at 8 minutes. The fluorescence intensity is presented as mean ± standard deviation (depicted in error bars) of two independent experiments. The maximum fluorescence intensity (100%) was derived from the positive control, Triton-X 0.1% (v/v). Fluorescence intensity (I) of the peptide-treated suspension was calculated and plotted as: (I_peptide_ / I_Triton-X(max)_) × 100%. The study was conducted as two independent experiments.

#### Molecular dynamics (MD) simulations

To understand the mode of interactions of the CaD23 peptide with the membranes, MD simulations of CaD23 with model bacterial and mammalian membranes were carried out. The distance between the centre of mass of CaD23 and the bilayer centre of mammalian and bacterial membranes is shown in **Figure 3A-B**. In the first 400 ns, the distance between CaD23 and both membranes decreased, suggesting a rapid adsorption of CaD23 on both membranes. CaD23 was closer to the bacterial membrane (z-distance = 2 nm) than the mammalian membrane (z-distance = 3.5 nm with considerable fluctuation), suggesting a stronger peptide-bacterial membrane interaction. Representative snapshots of the MD simulations of CaD23 with mammalian and bacterial membranes are shown in **Figure 4**.

**Figure 3.**
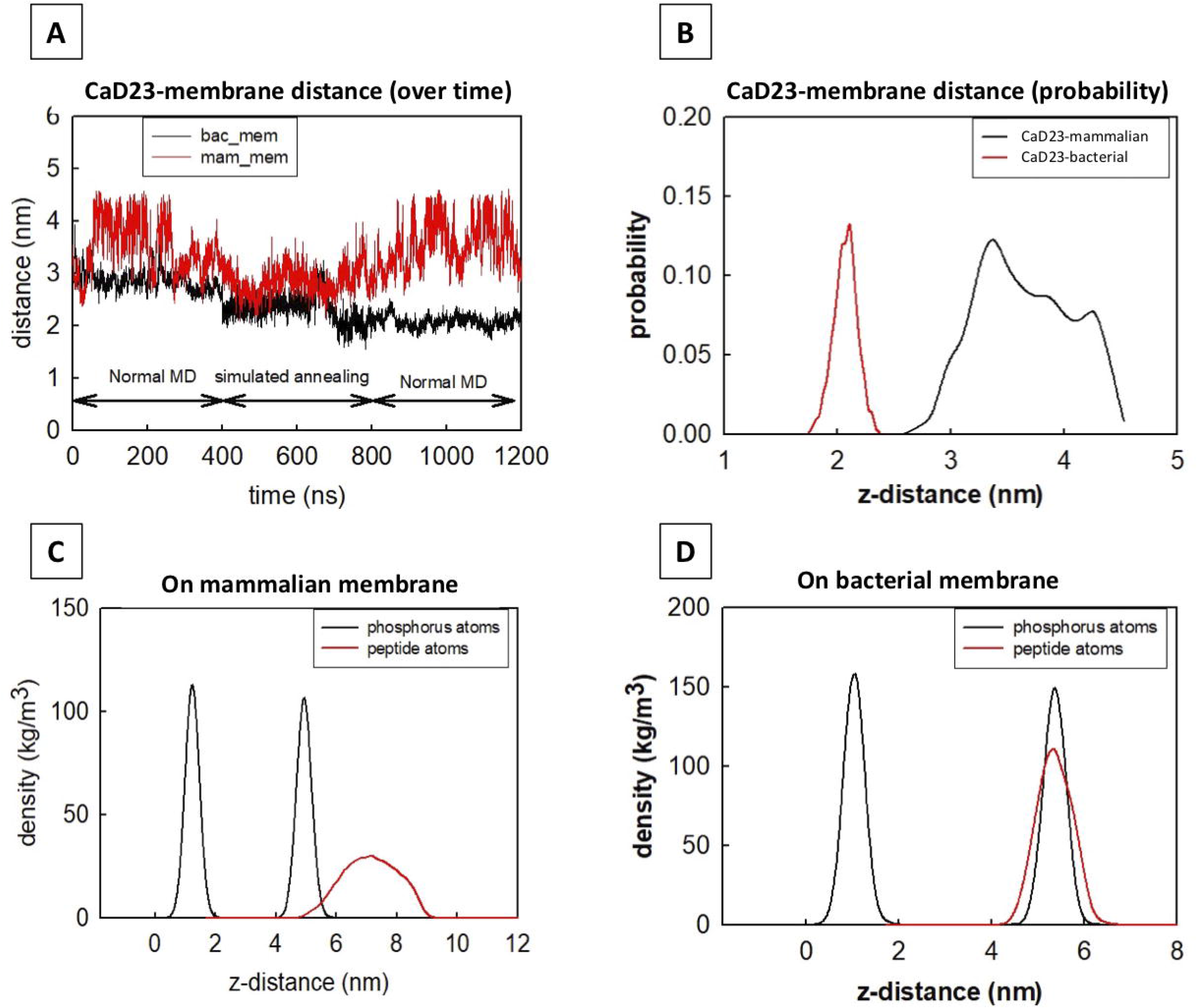
Molecular dynamics simulation of CaD23 on model mammalian and bacterial membranes. Each simulation was run for 400 ns at 300 K, followed by another 400 ns using simulated annealing (SA) to accelerate phase space sampling, finally followed by a further 400 ns simulation to obtain equilibration. **(A)** The graph showing the distance between CaD23 and mammalian or bacterial membrane over 1200 ns. CaD23 is shown to be closer to the bacterial membrane than to the mammalian membrane, suggesting a stronger interaction between CaD23 and the bacterial membrane. **(B)** The probability distribution of the peptide-membrane distance in the last 400 ns, demonstrating a closer distance of CaD23 to the bacterial membrane than to the mammalian membrane. **(C-D)** Density distributions of the CaD23 with respect to the phosphate groups of the bilayer membranes. The analysis is based on the last 400 ns simulation.

**Figure 4.**
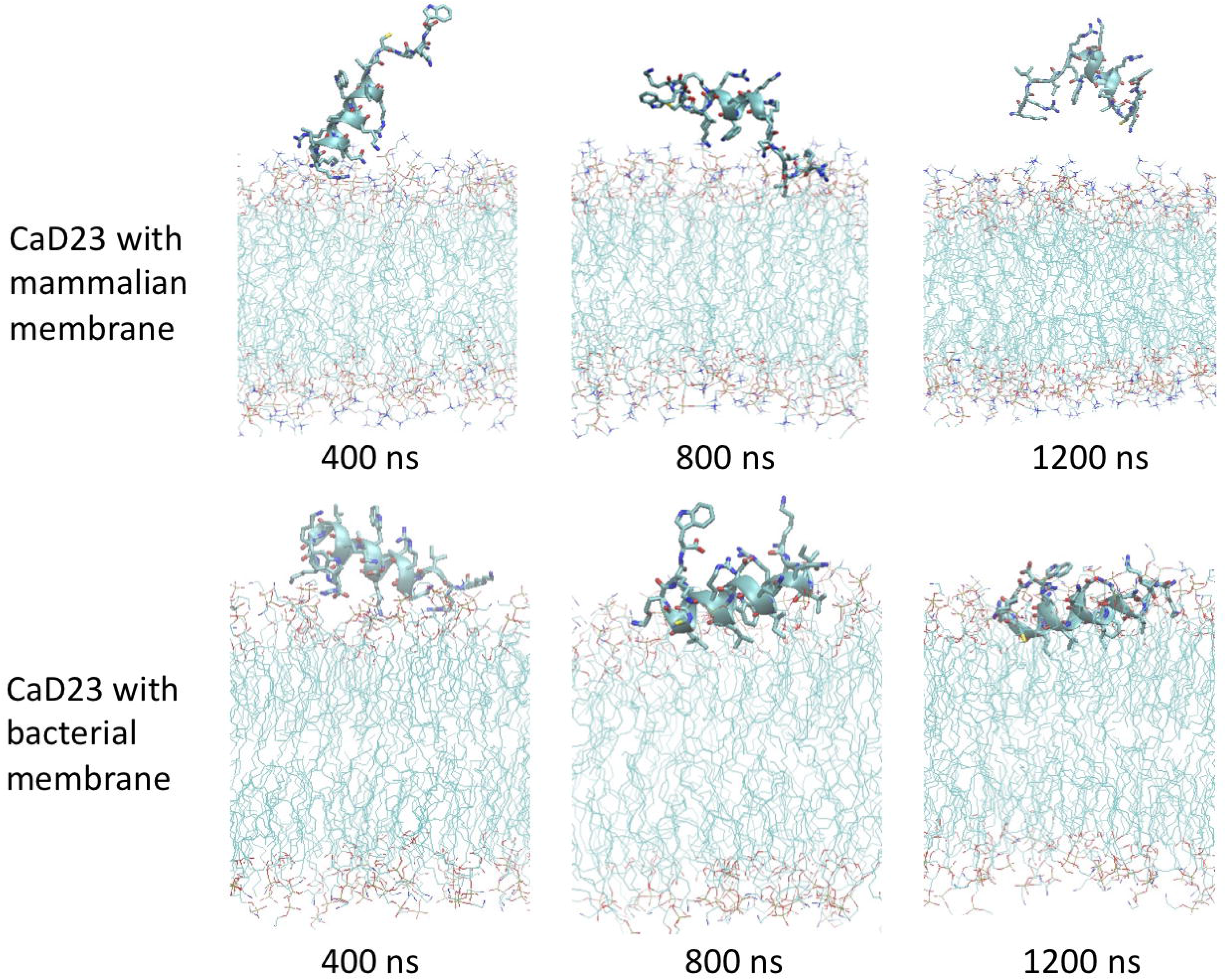
Molecular dynamics simulations study visualising the interaction between CaD23 and mammalian/bacterial membranes at an atomistic level. Representative snapshots of CaD23 with mammalian and bacterial membranes. The conformation of each snapshot corresponds to the most common configuration of CaD23 during the last 400 ns simulations. The snapshots demonstrate a stronger interaction (a closer distance) between CaD23 and bacterial membrane than mammalian membrane, corresponding with the experimental data. This also suggests that the rapid action of AMP23 is likely attributed to its membrane-permeabilising action.

Upon adsorption on the membrane, CaD23 started to interact with the head groups of the membrane, which involved the rearrangement of the head groups and the penetration of hydrophobic residues of CaD23 into the membrane (**Figure 4**). Due to complex mode of interactions, this process was characterized by a frustrated free energy landscape. To accelerate sampling, simulated annealing (SA) was applied. The peptide-membrane distance was found to decrease further, particularly for the distance between CaD23 and the bacterial membrane, because the strong perturbation of the bacterial head groups facilitated the penetration of the hydrophobic residues of CaD23 into the lipid tail region of the bacterial membrane, which did not occur on the mammalian membrane due to weak interactions. To obtain an equilibrium state, classical MD simulations without SA were carried out for a further 400 ns. The distance between the peptide and the bacterial membrane decreased further and remains stable. In contrast, the distance between CaD23 and the mammalian membrane increased and fluctuated with many adsorption-desorption events on the mammalian membrane, suggesting a weaker interaction.

The different locations of CaD23 with respect to the bilayer center can also be seen from the density distribution of CaD23 with respect to the phosphate atoms during the final 400 ns of the MD simulations (**Figure 3C-D**). On the mammalian membrane, the peak of CaD23 was low and the distribution of CaD23 was wide and far away from the phosphate groups, suggesting a low affinity of CaD23 to the mammalian membrane. In contrast, the peak of CaD23 was close to the phosphate groups upon strong adsorption on the bacterial membrane.

The helical wheel revealed that when the peptide was in helical conformation, it formed a perfect facial amphiphilic conformation, with positively charged residues facing one side and the hydrophobic residues facing the other side (**Figure 4**). Although the peptide largely maintained the helical conformation on both membranes, the peptide was more helical on the bacterial membrane than on the mammalian membrane. The snapshots from the last 400 ns in **Figure 4** clearly demonstrate that the peptide adopts a helical conformation on the bacterial membrane, with the hydrophobic residues inserted into the lipid tail region while the basic residues interact with the head groups, resulting in perturbation of the membrane-water interface. On the mammalian membrane, CaD23 was only partially helical and fluctuated due to the lack of strong electrostatic interactions, resulting in less perturbation of the mammalian membrane. Moreover, CaD23 formed more hydrogen bonds with the bacterial membrane compared to the mammalian membrane (**Figure 6**), which further contributed to the high affinity to the bacterial membrane.

**Figure 5.**
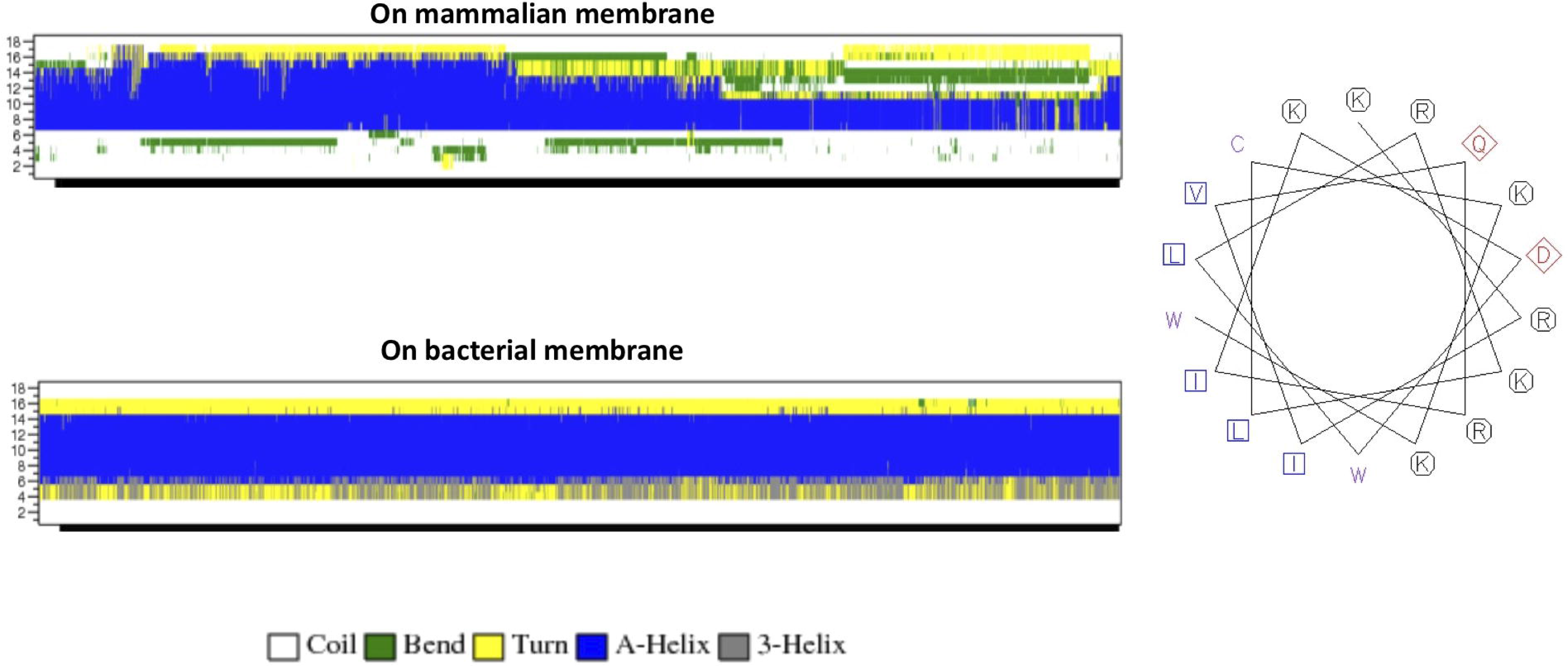
**(A)** Secondary structure evolution of CaD23 during the molecular dynamics (MD) simulations. CaD23 adopts a partially alpha-helical structure on the mammalian membrane compared to a highly alpha-helical structure on the bacterial membrane. **(B)** The helical wheel plot of CaD23. Blue and purple letters represent hydrophobic residues, red letters represent negatively charged acidic residues, and black letters represent positive charged basic residues.

**Figure 6.**
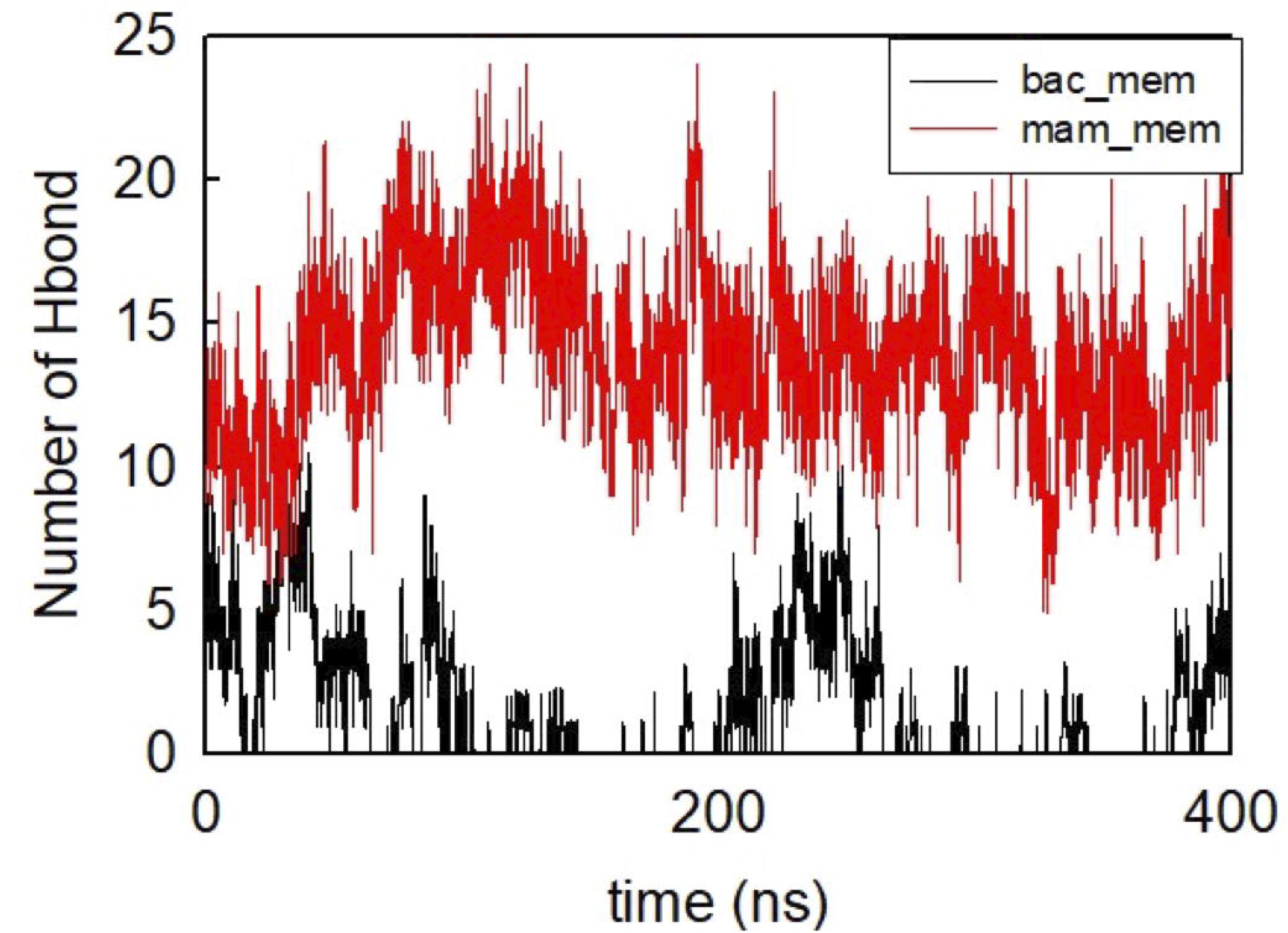
Number of hydrogen bond formed between CaD23 and the two membranes during the last 400 ns.

## DISCUSSION

The serendipitous discovery of HDPs in the early 1980s has sparked a significant interest in the field of antimicrobial therapy as HDPs have been shown to exhibit broad-spectrum and rapid antimicrobial action, with low risk of developing AMR. However, a number of barriers, including toxicity to host cells / tissues, have so far impeded the translation of HDP-based treatment to clinical use. In this study, we demonstrated that CaD23 could enhance the antimicrobial efficacy of commonly used antibiotics, including amikacin and levofloxacin, in a strong additive manner, against methicillin-sensitive and methicillin-resistant *S. aureus* when they were used in combination. This suggests that a lower treatment concentration of CaD23 and antibiotic can be used, serving as a useful strategy to reduce the concentration-dependent drug toxicity that is often observed in clinical practice.^48,49^

Furthermore, the addition of CaD23 at sub-MIC level was able to expedite the antimicrobial action of amikacin by 2-4 times when used in combination. Theoretically, such beneficial effect can reduce the risk of developing AMR as the bacteria have less time to adapt and develop effective mechanisms against the antibiotics. Studies have shown that membrane-active peptides with rapid antimicrobial action have a low risk of developing AMR whereas conventional antibiotics are prone to developing AMR, especially when they are chronically used at a sub-MIC level.^27,41,50^ This is due to the fact that modification of the entire membrane of the microorganisms in response to membrane-active peptides incurs a high fitness cost when compared to alteration of a particular binding site targeted by conventional antibiotics (e.g. alteration in the penicillin-binding protein reduces the efficacy of beta-lactam antibiotics).^29,51^

The advantageous strong additive effects of CaD23-amikacin and CaD23-levofloxacin against Gram-positive bacteria are likely attributed the different underlying mechanism of action of these drugs. Amikacin is a commonly used aminoglycoside in clinical practice (including ophthalmology) that exhibits its antimicrobial activity via inhibition of the 30S ribosomal subunit^6,52^ whereas levofloxacin, a frequently used fluoroquinolone, kills bacteria by inhibiting the bacterial DNA gyrase. It is likely that CaD23 interacts and permeabilises the cytoplasmic membrane of Gram-positive bacteria and facilitates the penetration of aminoglycoside and levofloxacin into the bacterial cells, enabling a more effective binding to the intracellular targets. However, none of the combinations exhibited an antagonistic effect, suggesting that both treatments display different mode of antimicrobial action.

Interestingly, we did not observe the same antimicrobial additive effect when CaD23 was used in combination with either amikacin or levofloxacin against Gram-negative bacteria. One of the main differences between Gram-positive and Gram-negative bacteria lies in the different compositions of the bacterial cell envelope.^53^ While both types of bacteria have a cytoplasmic / inner membrane, Gram-positive bacteria possess a thick peptidoglycan outer layer whereas Gram-negative bacteria possess an additional outer membrane, which is primarily composed of negatively charged lipopolysaccharides (in the outer leaflet of the outer membrane).^51^ It is likely that CaD23 primarily acts on the inner cell membrane (of both types of bacteria), with a weaker interaction with lipopolysaccharides, thereby explaining the additive effects of combined CaD23-antibiotic that were observed in Gram-positive bacteria but not in Gram-negative bacteria. Further investigations are warranted to understand the lack of mode of activity of CaD23-antibiotic combination against Gram-negative bacteria.

On the other hand, our group had recently demonstrated that FK16 (a truncated version of LL-37) was able to enhance the antimicrobial activity of vancomycin against *P. aeruginosa*.^32^ Vancomycin is a glycopeptide antibiotic that has poor permeability against the outer membrane of Gram-negative bacteria.^54^ It was hypothesised that FK16, a membrane-active peptide, permeabilizes the outer membrane of the Gram-negative bacteria and improves the delivery of vancomycin to access periplasmic cell wall precursors and intracellular target. Antonoplis et al.^54^ had similarly demonstrated the synergistic effect in a vancomycin-arginine peptide conjugate in treating carbapenem-resistant *Escherichia coli*, likely through a similar mechanism of action described above. Kampshoff et al.^31^ observed a synergistic effect between ciprofloxacin and melimine (a highly cationic, hybridised peptide derived from melittin and protamine) against ciprofloxacin-resistant *P. aeruginosa*, but not against *S. aureus* or non-drug resistant *P. aeruginosa*. In addition, a synergistic effect was not observed in either melimine-cefepime (a fourth-generation cephalosporin), Mel4 (truncated melimine)-ciprofloxacin, or Mel4-cefepim, highlighting the heterogeneous interactions among different types of peptides and antibiotics.

Both SYTOX green uptake assay and MD simulation studies demonstrated that CaD23 achieved its antimicrobial activity *via* a membrane-permeabilising action. In the recent decades, MD simulations have been increasingly utilised in the process of drug discovery and development in many fields, including the field of HDPs.^29,55–58^ They have been shown to predict the secondary structures of proteins / peptides, decipher the underlying mechanism of action, and identifying key residues responsible for the protein-protein or protein-membrane interaction at an atomistic level.^57,59,60^ As the chemical space of synthetic and natural HDPs is vast, MD simulation serves as a powerful tool to expedite the process of designing and optimising the peptide sequences as it reduces the need for repetitive microbiological assays and laborious screening of a large amount of peptide that is usually required in traditional mutation-based empirical methods.

A number of key factors, including alpha-helicity, amphiphilicity, cationicity and hydrophobicity, have been described to influence the antimicrobial efficacy of HDPs.^23,27,29^ In our study, MD simulations have revealed a number of important findings pertaining to the CaD23 molecule. Firstly, we observed a rapid adsorption of CaD23 on the negatively charged bacterial membrane during the early stage of the simulation (particularly at the N-terminus where the Lys1 is located), highlighting the importance of cationicity in the CaD23 molecule. In contrast, the zwitterionic nature of the mammalian membrane exhibited a weaker interaction with CaD23. Secondly, we showed that CaD23 adopted a more alpha-helical conformation on the bacterial membrane than the mammalian membrane, suggesting that alpha-helicity plays an important contributory role to the antimicrobial efficacy of CaD23. In the helical conformation, the peptide displays high facial amphiphilicity, which resulted in a more favourable interaction with the bacterial membrane, with a deeper penetration of CaD23 into the bacterial membrane. This is in accordance with many studies in the literature that had highlighted the important correlation between alpha-helicity and antimicrobial efficacy observed in various natural and synthetic HDPs.^23,26^ We also observed that the Trp18 residue at the C-terminal had a strong interaction with the bacterial membrane but not the Trp10 residue. This suggests that the Trp10 residue may potentially be substituted with a less hydrophobic residue such as Leu or Ile to reduce the hydrophobicity and toxicity, and to improve its water solubility.

Despite the many advantages of MD simulations described above, it is noteworthy to mention that the model bacterial membrane utilised in the current MD simulation is only representative of the inner membrane of the Gram-positive and Gram-negative bacteria. Atomistic models have been developed for bacterial outer membrane and several studies have been carried out to understand the structural dynamics of the outer membrane.^61–63^ However, MD simulations with outer membrane is out of the scope of this study as CaD23 was mainly efficacious against Gram-positive bacteria.

In summary, our study demonstrated that CaD23 is a membrane-active peptide that has the ability to enhance the antimicrobial action of commonly used antibiotics such as amikacin and levofloxacin, potentially offering a new therapeutic strategy for Gram-positive bacterial infection. Further *in vivo* studies to validate these results would be invaluable. In addition, MD simulation serves as a useful computational tool in deciphering the underlying mechanism of action and guiding the design process of HDPs.

## AUTHOR CONTRIBUTION STATEMENT

Study conceptualisation and design: DSJT, IM, HSD

Data collection: DSJT, JL, ETLG

Data analysis and interpretation: DSJT, JL, CSV, DGS, MN, LM, RWB, RL, IM, HSD

Manuscript drafting: DSJT, JL

Critical revision of manuscript: CSV, ETLG, DGS, MN, LM, RWB, RL, IM, HSD

Final approval of manuscript: All authors

## REFERENCES

1. Prestinaci F, Pezzotti P, Pantosti A. Antimicrobial resistance: a global multifaceted phenomenon. Pathog Glob Health. 2015;109(7):309–18.

2. Ventola CL. The antibiotic resistance crisis: part 1: causes and threats. P t. 2015;40(4):277–83.

3. O’Neill J. Tackling drug-resistant infections globally: Final report and recommendations. Review on Antimicrobial Resistance. 2016:1–81.

4. Ting DSJ, Ho CS, Deshmukh R, Said DG, Dua HS. Infectious keratitis: an update on epidemiology, causative microorganisms, risk factors, and antimicrobial resistance. Eye (Lond). 2021;35(4):1084–101.

5. Pulido-Cejudo A, Guzmán-Gutierrez M, Jalife-Montaño A, Ortiz-Covarrubias A, Martínez-Ordaz JL, Noyola-Villalobos HF, et al. Management of acute bacterial skin and skin structure infections with a focus on patients at high risk of treatment failure. Ther Adv Infect Dis. 2017;4(5):143–61.

6. Ting DSJ, Ho CS, Cairns J, Elsahn A, Al-Aqaba M, Boswell T, et al. 12-year analysis of incidence, microbiological profiles and in vitro antimicrobial susceptibility of infectious keratitis: the Nottingham Infectious Keratitis Study. Br J Ophthalmol. 2021;105(3):328–33.

7. Hoffman JJ, Burton MJ, Leck A. Mycotic Keratitis-A Global Threat from the Filamentous Fungi. J Fungi (Basel). 2021;7(4).

8. Khoo P, Cabrera-Aguas MP, Nguyen V, Lahra MM, Watson SL. Microbial keratitis in Sydney, Australia: risk factors, patient outcomes, and seasonal variation. Graefes Arch Clin Exp Ophthalmol. 2020;258(8):1745–55.

9. Ting DSJ, Ghosh N, Ghosh S. Herpes zoster ophthalmicus. BMJ. 2019;364:k5234.

10. Ting DSJ, Settle C, Morgan SJ, Baylis O, Ghosh S. A 10-year analysis of microbiological profiles of microbial keratitis: the North East England Study. Eye (Lond). 2018;32(8):1416–7.

11. Green M, Carnt N, Apel A, Stapleton F. Queensland Microbial Keratitis Database: 2005-2015. Br J Ophthalmol. 2019;103(10):1481–6.

12. Khor WB, Prajna VN, Garg P, Mehta JS, Xie L, Liu Z, et al. The Asia Cornea Society Infectious Keratitis Study: A Prospective Multicenter Study of Infectious Keratitis in Asia. Am J Ophthalmol. 2018;195:161–70.

13. Shah A, Sachdev A, Coggon D, Hossain P. Geographic variations in microbial keratitis: an analysis of the peer-reviewed literature. Br J Ophthalmol. 2011;95(6):762–7.

14. Ting DSJ, Bignardi G, Koerner R, Irion LD, Johnson E, Morgan SJ, et al. Polymicrobial Keratitis With Cryptococcus curvatus, Candida parapsilosis, and Stenotrophomonas maltophilia After Penetrating Keratoplasty: A Rare Case Report With Literature Review. Eye Contact Lens. 2019;45(2):e5–e10.

15. Tu EY, Joslin CE, Nijm LM, Feder RS, Jain S, Shoff ME. Polymicrobial keratitis: Acanthamoeba and infectious crystalline keratopathy. Am J Ophthalmol. 2009;148(1):13–9.e2.

16. Asbell PA, Sanfilippo CM, Sahm DF, DeCory HH. Trends in Antibiotic Resistance Among Ocular Microorganisms in the United States From 2009 to 2018. JAMA Ophthalmol. 2020;138(5):439–50.

17. Hernandez-Camarena JC, Graue-Hernandez EO, Ortiz-Casas M, Ramirez-Miranda A, Navas A, Pedro-Aguilar L, et al. Trends in Microbiological and Antibiotic Sensitivity Patterns in Infectious Keratitis: 10-Year Experience in Mexico City. Cornea. 2015;34(7):778–85.

18. Lalitha P, Manoharan G, Karpagam R, Prajna NV, Srinivasan M, Mascarenhas J, et al. Trends in antibiotic resistance in bacterial keratitis isolates from South India. Br J Ophthalmol. 2017;101(2):108–13.

19. Lin L, Duan F, Yang Y, Lou B, Liang L, Lin X. Nine-year analysis of isolated pathogens and antibiotic susceptibilities of microbial keratitis from a large referral eye center in southern China. Infect Drug Resist. 2019;12:1295–302.

20. Ting DSJ, Henein C, Said DG, Dua HS. Photoactivated chromophore for infectious keratitis-corneal cross-linking (PACK-CXL): A systematic review and meta-analysis. Ocul Surf. 2019;17(4):624–34.

21. Ting DSJ, Henein C, Said DG, Dua HS. Amniotic membrane transplantation for infectious keratitis: a systematic review and meta-analysis. Sci Rep. 2021;11(1):13007.

22. Hossain P, Tourkmani AK, Kazakos D, Jones M, Anderson D. Emergency corneal grafting in the UK: a 6-year analysis of the UK Transplant Registry. Br J Ophthalmol. 2018;102(1):26–30.

23. Mookherjee N, Anderson MA, Haagsman HP, Davidson DJ. Antimicrobial host defence peptides: functions and clinical potential. Nat Rev Drug Discov. 2020.

24. Mohammed I, Said DG, Dua HS. Human antimicrobial peptides in ocular surface defense. Prog Retin Eye Res. 2017;61:1–22.

25. Hancock RE, Lehrer R. Cationic peptides: a new source of antibiotics. Trends Biotechnol. 1998;16(2):82–8.

26. Haney EF, Straus SK, Hancock REW. Reassessing the Host Defense Peptide Landscape. Front Chem. 2019;7:43.

27. Hancock RE, Sahl HG. Antimicrobial and host-defense peptides as new anti-infective therapeutic strategies. Nat Biotechnol. 2006;24(12):1551–7.

28. Hancock RE, Haney EF, Gill EE. The immunology of host defence peptides: beyond antimicrobial activity. Nature reviews Immunology. 2016;16(5):321–34.

29. Ting DSJ, Beuerman RW, Dua HS, Lakshminarayanan R, Mohammed I. Strategies in Translating the Therapeutic Potentials of Host Defense Peptides. Front Immunol. 2020;11:983.

30. Lakshminarayanan R, Tan WX, Aung TT, Goh ET, Muruganantham N, Li J, et al. Branched peptide, B2088, disrupts the supramolecular organization of lipopolysaccharides and sensitizes the Gram-negative bacteria. Sci Rep. 2016;6:25905.

31. Kampshoff F, Willcox MDP, Dutta D. A Pilot Study of the Synergy between Two Antimicrobial Peptides and Two Common Antibiotics. Antibiotics (Basel). 2019;8(2).

32. Mohammed I, Said DG, Nubile M, Mastropasqua L, Dua HS. Cathelicidin-Derived Synthetic Peptide Improves Therapeutic Potential of Vancomycin Against Pseudomonas aeruginosa. Front Microbiol. 2019;10:2190.

33. Nuding S, Frasch T, Schaller M, Stange EF, Zabel LT. Synergistic effects of antimicrobial peptides and antibiotics against Clostridium difficile. Antimicrob Agents Chemother. 2014;58(10):5719–25.

34. Pletzer D, Mansour SC, Hancock REW. Synergy between conventional antibiotics and anti-biofilm peptides in a murine, sub-cutaneous abscess model caused by recalcitrant ESKAPE pathogens. PLoS Pathog. 2018;14(6):e1007084.

35. Mishra B, Reiling S, Zarena D, Wang G. Host defense antimicrobial peptides as antibiotics: design and application strategies. Curr Opin Chem Biol. 2017;38:87–96.

36. Ting DSJ, Goh ETL, Mayandi V, Busoy JMF, Aung TT, Periayah MH, et al. Hybrid Derivative of Cathelicidin and Human Beta Defensin-2 Against Gram-Positive Bacteria: A Novel Approach for the Treatment of Bacterial Keratitis. bioRxiv. 2021: 2021.04.22.440925.

37. Ting DSJ, Cairns J, Gopal BP, Ho CS, Krstic L, Elsahn A, et al. Risk Factors, Clinical Outcomes and Prognostic Factors of Bacterial Keratitis: The Nottingham Infectious Keratitis Study. medRxiv. 2021:2021.05.26.21257881.

38. Borkar DS, Fleiszig SM, Leong C, Lalitha P, Srinivasan M, Ghanekar AA, et al. Association between cytotoxic and invasive Pseudomonas aeruginosa and clinical outcomes in bacterial keratitis. JAMA Ophthalmol. 2013;131(2):147–53.

39. Lee EJ, Truong TN, Mendoza MN, Fleiszig SM. A comparison of invasive and cytotoxic Pseudomonas aeruginosa strain-induced corneal disease responses to therapeutics. Curr Eye Res. 2003;27(5):289–99.

40. Clinical & Laboratory Standards Institute (CLSI). Methods for dilution antimicrobial susceptibility tests for bactera that grow aerobically, 11th edition. 2019.

41. Mayandi V, Xi Q, Leng Goh ET, Koh SK, Jie Toh TY, Barathi VA, et al. Rational Substitution of ε-Lysine for α-Lysine Enhances the Cell and Membrane Selectivity of Pore-Forming Melittin. J Med Chem. 2020;63(7):3522–37.

42. Jorgensen WL, Chandrasekhar J, Madura JD, Impey RW, Klein ML. Comparison of simple potential functions for simulating liquid water. J Chem Phys. 1983;79(2):926–35.

43. Farrotti A, Bocchinfuso G, Palleschi A, Rosato N, Salnikov ES, Voievoda N, et al. Molecular dynamics methods to predict peptide locations in membranes: LAH4 as a stringent test case. Biochim Biophys Acta. 2015;1848(2):581–92.

44. Hess B, Bekker H, Berendsen HJC, Fraaije JGE. LINCS: A linear constraint solver for molecular simulations. J Computational Chemistry. 1998;18(12):1463–72.

45. Essmann U, Perera L, Berkowitz ML. A smooth particle mesh Ewald method. J Chem Phys. 1995;103(19):8577.

46. Nose S, Klein ML. Constant pressure molecular dynamics for molecular systems. Mol Phys. 1983;50(5):1055–76.

47. Martonák R, Laio A, Parrinello M. Predicting crystal structures: the Parrinello-Rahman method revisited. Phys Rev Lett. 2003;90(7):075503.

48. Dua HS, Otri AM, Said DG, Faraj LA. The ‘up-down’ sign of acute ocular surface drug toxicity. Br J Ophthalmol. 2012;96(11):1439–40.

49. Forster AJ, Murff HJ, Peterson JF, Gandhi TK, Bates DW. Adverse drug events occurring following hospital discharge. J Gen Intern Med. 2005;20(4):317–23.

50. Llor C, Bjerrum L. Antimicrobial resistance: risk associated with antibiotic overuse and initiatives to reduce the problem. Ther Adv Drug Saf. 2014;5(6):229–41.

51. Kapoor G, Saigal S, Elongavan A. Action and resistance mechanisms of antibiotics: A guide for clinicians. J Anaesthesiol Clin Pharmacol. 2017;33(3):300–5.

52. Krause KM, Serio AW, Kane TR, Connolly LE. Aminoglycosides: An Overview. Cold Spring Harb Perspect Med. 2016;6(6).

53. Silhavy TJ, Kahne D, Walker S. The bacterial cell envelope. Cold Spring Harb Perspect Biol. 2010;2(5):a000414.

54. Antonoplis A, Zang X, Wegner T, Wender PA, Cegelski L. Vancomycin-Arginine Conjugate Inhibits Growth of Carbapenem-Resistant E. coli and Targets Cell-Wall Synthesis. ACS Chem Biol. 2019;14(9):2065–70.

55. Durrant JD, McCammon JA. Molecular dynamics simulations and drug discovery. BMC Biol. 2011;9:71.

56. De Vivo M, Masetti M, Bottegoni G, Cavalli A. Role of Molecular Dynamics and Related Methods in Drug Discovery. J Med Chem. 2016;59(9):4035–61.

57. Li J, Koh JJ, Liu S, Lakshminarayanan R, Verma CS, Beuerman RW. Membrane Active Antimicrobial Peptides: Translating Mechanistic Insights to Design. Front Neurosci. 2017;11:73.

58. Li J, Lakshminarayanan R, Bai Y, Liu S, Zhou L, Pervushin K, et al. Molecular dynamics simulations of a new branched antimicrobial peptide: a comparison of force fields. J Chem Phys. 2012;137(21):215101.

59. Li J, Beuerman RW, Verma CS. Molecular Insights into the Membrane Affinities of Model Hydrophobes. ACS Omega. 2018;3(3):2498–507.

60. Tsai CW, Hsu NY, Wang CH, Lu CY, Chang Y, Tsai HH, et al. Coupling molecular dynamics simulations with experiments for the rational design of indolicidin-analogous antimicrobial peptides. J Mol Biol. 2009;392(3):837–54.

61. Pontes FJ, Rusu VH, Soares TA, Lins RD. The Effect of Temperature, Cations, and Number of Acyl Chains on the Lamellar to Non-Lamellar Transition in Lipid-A Membranes: A Microscopic View. J Chem Theory Comput. 2012;8(10):3830–8.

62. Li J, Beuerman R, Verma CS. Dissecting the Molecular Mechanism of Colistin Resistance in mcr-1 Bacteria. J Chem Inf Model. 2020;60(10):4975–84.

63. Li J, Beuerman R, Verma CS. Mechanism of polyamine induced colistin resistance through electrostatic networks on bacterial outer membranes. Biochim Biophys Acta Biomembr. 2020;1862(9):183297.

