## Supplementary material for "Evaluation of Host Defense Peptide (CaD23)-Antibiotic Interaction and Mechanism of Action: Insights from Experimental and Molecular Dynamics Simulations Studies": Table 1

**Table 1.** Minimum inhibitory concentration (MIC) of CaD23, amikacin and levofloxacin against methicillin-sensitive *Staphylococcus aureus* (SH1000), methicillin-resistant *S. aureus* (ATCC MRSA43300), *Pseudomonas aeruginosa* ATCC PA19660 (cytotoxic strain), ATCC PA27853 (invasive strain) and PAO1L (invasive strain). The MIC value is expressed in μg/ml (and μM in bracket).

| **Treatment** | **SH1000** | **MRSA43300** | **PA19660** | **PA27853** | **PAO1L** |
| --- | --- | --- | --- | --- | --- |
| CaD23 | 12.5 (5.2) | 25 (10.4) | 25 (10.4) | 25 (10.4) | 50 (20.8) |
| Amikacin | 1.25 (2.1) | 2.5 (4.3) | 0.63 (1.1) | 1.25 (2.1) | 0.63 (1.1) |
| Levofloxacin | 0.31 (0.86) | 0.31 (0.86) | 0.31 (0.86) | 0.63 (1.7) | 0.31 (0.86) |

All assays were conducted as two independent experiments in biological duplicate.
